# Motion tolerance in wearable OPM-MEG using dynamic field nulling

**DOI:** 10.64898/2026.08.17.745285

**Authors:** Mainak Jas, Teppei Matsubara, Steven M. Stufflebeam, Padmavathi Sundaram, Seppo P. Ahlfors

## Abstract

Wearable magnetoencephalography (MEG) enabled by optically pumped magnetometers (OPMs) promises improved comfort and motion tolerance. This is particularly beneficial when measuring brain activity in children who cannot sit still for long periods of time. Compared to cryogenic MEG, wearable MEG allows larger head movements, but they result in artifacts due to uncompensated background fields and reduce source localization accuracy.

Spatial filtering methods can partially compensate these motion-induced artifacts, but they are most effective when used in combination with background field nulling. This is because accurate spatial filtering relies on an accurate estimate of the sensor gain and orientation of its sensitive axis. Through simulations, we first deduce the target residual background field that is necessary for accurate dipole localization (< 1 cm) in the presence of head movements. Our simulations indicate that smaller background fields enable larger head movements. Then, we used our open-source printed circuit board (PCB) coils to develop a method to dynamically null the background field. We demonstrate that our dynamic field nulling method allows improved localization of somatosensory evoked fields (SEFs) by maintaining the background field below the target residual fields established in the simulations. Our study highlights the importance of tracking both the background field and the head position relative to the background field for quality assurance in wearable MEG.

## 1. Introduction

Magnetoencephalography (MEG) is an established non-invasive tool for measuring neural activity with high temporal resolution (M. Hämäläinen et al., 1993a). Conventional MEG systems use cryogenic sensors within a rigid gantry and require participants to remain stationary during measurements, limiting their application in naturalistic settings and in patient cohorts where motion is inevitable. For example, in children with epilepsy, conventional MEG is not routinely used despite offering improved diagnostic yield over using EEG alone (Geller et al., 2023). Addressing this limitation, a new wearable MEG sensor technology based on optically pumped magnetometers (OPMs) is emerging that enables natural motion during recordings and offers greater comfort and tolerability amongst pediatric populations (Boto et al., 2018).

However, achieving true motion tolerance in wearable MEG remains a significant technical challenge. Unlike cryogenic systems where relatively small head movements occur within a fixed sensor array, wearable MEG allows the sensors themselves to move within a background field. Even very small background fields are sufficient to induce artifacts that can contaminate the data and distort the source localization accuracy of MEG. Head movement changes the projection of the field onto the sensor’s sensitive axes, generating motion-induced artifacts. These artifacts can arise from rotation within a uniform field, or from linear and rotational movement within a field gradient.

Previous work has characterized how background magnetic fields influence the performance of OPMs, including their effects on sensor non-linearity and sensitivity (Borna et al., 2020; Nardelli et al., 2019). Mitigation strategies combine hardware and software approaches including active field nulling (Holmes et al., 2018, 2019; Rea et al., 2021) and post hoc correction using spatial filtering methods (M. S. Hämäläinen & Ilmoniemi, 1994; McPherson et al., 2025; Tierney et al., 2021, 2024; Yu et al., 2026). Nevertheless, the relationship between head motion, residual fields, and localization accuracy remains poorly quantified.

This question is particularly relevant in epilepsy, where individual transient events are clinically meaningful (Murakami et al., 2016), so even small motion-induced errors can bias localization. For instance, interictal epileptiform discharges (IEDs) localized away from the cluster center of other IEDs may be misinterpreted as evidence of propagation or an extended epileptogenic zone. In wearable OPM-MEG, this apparent propagation could instead reflect motion artifacts arising from head movements within the residual background field. Thus, clear clinical criteria are needed on the allowable background field for varying amounts of head motion, to reliably distinguish apparent propagation from true propagation of IEDs.

While hardware-based background field suppression is common, it remains unknown whether the residual field obtained is sufficient for clinically acceptable localization accuracy (< 1 cm), for a given amount of motion. Here, we address this problem using a combination of simulations and hardware development. First, we quantify how background field and head motion jointly affect dipole localization accuracy and compare this to conventional cryogenic MEG. We show that cross-axis projection error (CAPE) due to residual background fields can reduce the effectiveness of spatial filtering methods in reducing motion-induced artifacts, and that this holds true even with no field gradients using only rotational motion within a uniform background field. Such rotational motion is common in natural head movements and is abundantly observed in children with epilepsy. Second, we present a dynamic field nulling approach based on our previously published open-source printed circuit board (PCB) coil system (Jas et al., 2025), using orthogonally arranged reference sensors and proportional-integral (PI) control to suppress uniform background fields. This provides continuous monitoring and reduction of background fields during acquisition. Finally, we show that dynamic field nulling can reduce background fields and improve motion robustness. We share our hardware and software design to support adoption and further development. Together, this work provides quantitative guidance on motion tolerance in wearable MEG, along with practical tools to improve performance in clinical and real-world settings.

## 2. Methods

### 2.1. Understanding motion artifacts in wearable MEG

**Figure 1.**
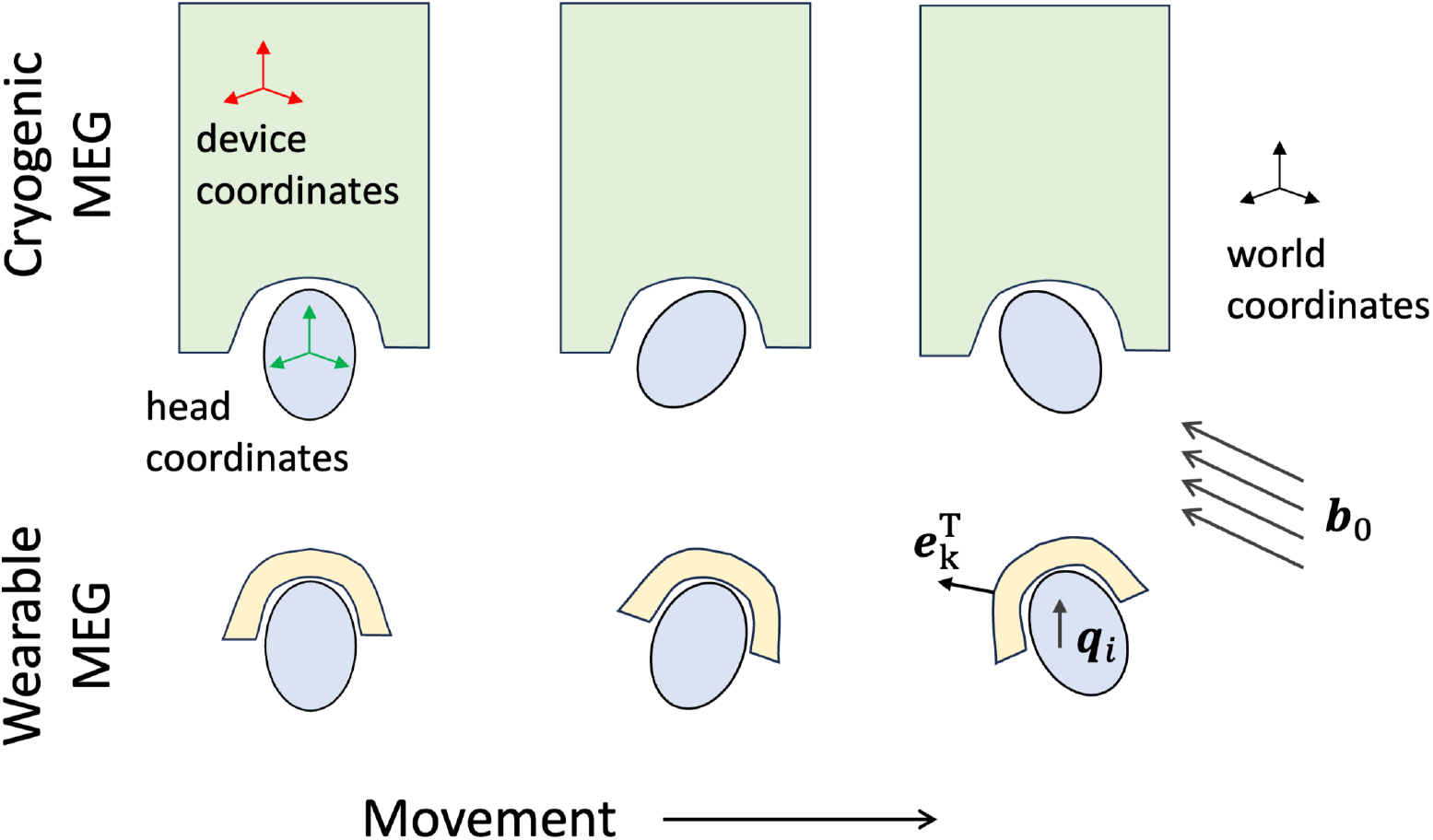
Conceptual figure depicting the origin of motion artifacts in cryogenic *vs*. wearable MEG. While motion artifacts in cryogenic MEG are due to changes in the relative sensitivity of magnetometers to brain activity, the artifacts in wearable MEG are due to changes in the relative sensitivity to external magnetic field.

Movement artifacts in cryogenic MEG and wearable MEG arise due to different reasons. In cryogenic MEG, the device is fixed inside the magnetically shielded room, but the head can move inside the dewar, particularly with pediatric head sizes. While this head movement is not significant, it can change the relative strength of the magnetic field detected by external magnetometers. Contrasting this to wearable MEG, the head is static with respect to the sensor helmet, but the sensors move with respect to the background field. As the sensors move within the background field, they detect changes in the sensed field which causes movement artifacts.

We can model these differences mathematically. Both systems measure the summation of background fields **b**_0_ ∈ ℝ^3^ and brain activity, but the time varying motion component impacts the measured magnetic field differently. In a cryogenic MEG system, we have

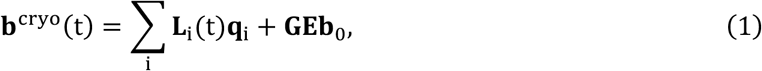

where **b**^cryo^(t) ∈ **R**^S^ is the measured magnetic field, **L**_i_(t) ∈ ℝ^S×3^ is the continuously varying lead field of the dipole **q**_i_ ∈ ℝ^3^ as the head moves within the fixed array of *S* sensors. The sensor gains **G** = diag(*g*_1_, …, *g*_*S*_) measures the sensor gains and **E** ∈ ℝ^S×3^ contains the unit vector in the direction of the sensors’ sensitive axes 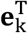.

In a wearable magnetometer, the lead field **L**_i_ is fixed in time because the helmet is strapped to the participant, but the sensor orientation changes relative to the background field:

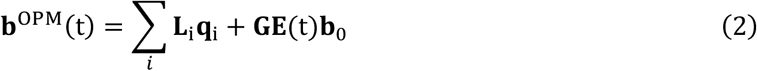

### 2.2. Compensation of motion artifacts

Similar to cryogenic MEG, motion artifacts can be compensated using spatial filtering family of methods such as Homogeneous Field Compensation (HFC) (Tierney et al., 2021, 2024). Since the measured artifact in Equation (2) is governed by a linear relationship, the background field can be estimated as a spatial pattern and a projector can be constructed to remove it. The artifacts measured across all sensors can be modeled as:

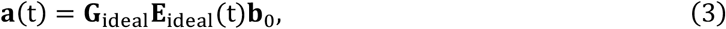

where **E**_ideal_ are the calibrated ideal sensitive axes. Spatial filtering methods builds a projector **P** onto the column space of **G**_ideal_**E**_ideal_(t) that removes any data component lying in it such that (**I** ™ **P**)**a**(t) = **0**.

This compensation assumes that the sensor’s true sensitive axis and gain match the ideal calibrated values during the initial field zeroing. In a zero-field optically pumped magnetometer (OPM) that operate in the spin-exchange-relaxation-free (SERF) regime, cross-axis projection error (CAPE) is a miscalibration that is introduced by transverse magnetic fields, tilting its sensitive axis 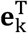 and changing its gain *g*_*k*_ (Borna et al., 2022). With a remnant transverse field *b*_*z*0_ at sensor *k*, the true axis is rotated by

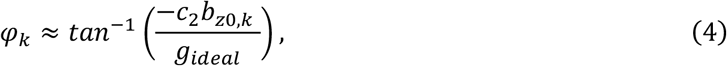

and the gain becomes

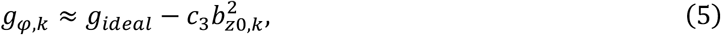

with sensor-specific coefficients *c*_2_ and *c*_3_ determined by the optical pumping rate, relaxation time, and modulation parameters. The rotation of the sensitive axis is frequency-dependent and has been estimated to be around 3^°^/nT of remnant field at 25 Hz. With CAPE, the remnant artifact after the projection becomes

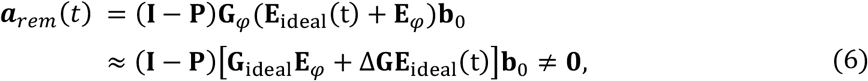

where **E**_φ_ ∈ ℝ^S×3^ contains the sensitive axis tilts, **G**_φ_ = diag(*g*_*φ*,1_, …, *g*_*φ,S*_), and Δ**G** = **G**_*φ*_ ™ **G**_ideal_. While CAPE degrades source localization accuracy even when stationary, head movement amplifies the impact of CAPE with higher amplitude movement artifact. Compensation based on spatial filtering only partially removes head movement artifacts in the presence of CAPE, leading to dipole localization errors.

### 2.3. Simulation of motion in uniform fields

To understand the relationship between head movement and background fields, we simulated the impact of head movement on the dipole localization accuracy.

Since we do not have access to head movement trajectories in wearable OPM-MEG, we used the 15 seconds of continuous head position indicator (cHPI) data from the MNE sample testing data and scaled the movement trajectories to obtain plausible movement patterns with wearable MEG. We simulated the impact of head movement on the estimated dipole localization accuracy in both cryogenic MEG and wearable OPM-MEG. The source time course was simulated with five randomly occurring interictal spikes (difference of Gaussians with amplitudes 300 nAm and 120 nAm) at the center of the ‘superior temporal’ label of the ‘aparc’ Freesurfer parcellation. We simulated three additional motion trajectories with rotational movements scaled by factors of 0.5, 2, and 4. The maximum rotation was 9.4° in the original motion trajectory. Errors in sensor gain and tilt due to CAPE were estimated by numerically solving the Bloch equations based on code provided in (Borna et al., 2022). Single-axis OPM sensors were created by projecting the SQUID magnetometers to be 5 mm above the scalp surface with their sensitive axis oriented normal to the scalp. For simulations with different number of sensors, the sensors closest to the source (as measured by Euclidean distance) were chosen. To simulate the maximum effect of CAPE, the dominant direction of sensor orientations was determined using a singular value decomposition (SVD) and the background field was oriented transverse to this direction. Since the head position is monitored less frequently than the sampling rate, the MEG data was interpolated in the intervening time points. The measured noise was assumed to be additive and comprising the intrinsic sensor noise (2.5 *fT*/√*Hz* for SQUID and 15 *fT*/√*Hz* for OPM), the background brain activity, and external non-brain background activity. We estimated the background brain activity noise from noise covariance in SQUID-MEG (in the sample dataset), projected it to source space using the SQUID-MEG inverse model, and projected it back to the OPM-MEG sensor space using the corresponding forward model. After filtering the data between 1 and 100 Hz, dipole estimation was performed at the peak of the interictal spike, and the localization error was computed as the Euclidean distance from the simulated dipole location. The dipole localization errors were averaged across N=10 simulations, each initialized with a different random seed.

For the base simulation, we assumed a background field of 1 nT and compared the dipole localization accuracy in wearable OPM-MEG (102 sensors) with cryogenic MEG for different amounts of head movement. Next, we varied the background field from 0 to 5 nT at intervals of 0.6 nT. After finding the interval in which the 1 cm dipole localization accuracy was achieved, the background field threshold estimate was refined by sampling every 0.06 nT within that interval. We repeated this process for each of the motion trajectories and for sensor arrays with 32, 64, and 102 sensors.

### 2.4. Background field nulling for minimizing motion artifacts

Background field nulling can mitigate motion artifacts by reducing the magnitude of the background field in a target volume (**Fig 2A**). In our prior work (Jas et al., 2025), we developed a PCB-based field nulling system for removing uniform and gradient components of the background fields allowing up to 20 cm head movements within a spherical volume of 50 cm diameter. Our measurements were performed with 18 OPMs (Gen 1, Fieldline, Colorado, USA) out of which 3 sensors were designated as reference sensors. These sensors were placed inside an 3D printed cast for measuring the orthogonal field components in the x (back of shielded room to front), y (floor to ceiling), and z (ear to ear) directions. A low-noise programmable current driver supplying up to 50 mA current (CSB-50 Twinleaf, Princeton, USA) is used for field nulling. We consider three different approaches for field nulling and tested their impact on data quality:

**Figure 2A.**
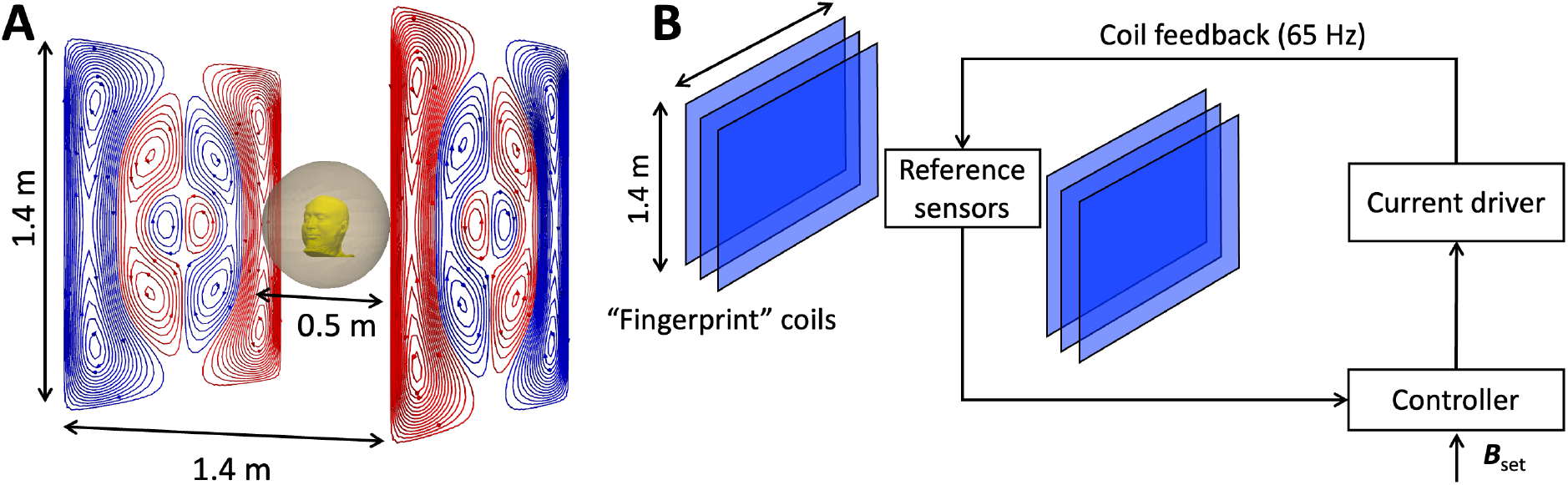
PCB-based dynamic field nulling removes background field in a 50 cm volume allowing up to 20 cm head movement within the target region depending on head size and positioning. Figure drawn to scale. **B**. Schematic of dynamic field nulling using a PCB-based nulling coils.

#### No field nulling

In this condition, the sensors were operated in closed-loop mode, but no additional external field nulling was applied. As newer OPMs can cancel up to 150 nT background field with a dynamic range of 30 nT (Alem et al., 2023), it is possible to perform wearable MEG measurements even with large background fields. This condition provides a baseline measurement for the data quality. When operated in this mode, cross-axis projection error (CAPE) is minimized (Borna et al., 2022). However, movement artifacts are still expected due to the remnant field in the region around the subject’s head which has to be compensated through postprocessing.

#### Static field nulling

After tuning the currents driving the coils to achieve near-zero field (< 2 nT) at the beginning of the experiment, the currents were kept fixed through the rest of the measurement. The tuning was performed using an iterative method where the coil current was updated with a step size equal to the inverse of the coil-to-sensor sensitivity (*B*_*x*_ = 0.68 mA/nT, *B*_*y*_= 0.62 mA/nT, *B*_*z*_ = 0.14 mA/nT). The iterations continued until the residual background field, as measured by the on-sensor coils after field coarse zeroing was less than 2 nT. A final fine zeroing step was performed to get an accurate estimate of the residual field. In this mode, the movement artifacts were expected to be reduced compared to the condition where no PCB-based field nulling is applied. The sensors were operated in closed loop after the initial static field nulling to minimize CAPE even if the field drifted from the initial values.

#### Dynamic field nulling

In the method of dynamic field nulling, continuous adjustment of the background field was achieved using a closed-loop proportional integral (PI) control (**Fig 2B**). Before dynamic field nulling, static field nulling was performed to prevent the on-sensor coils from saturating. After the initial static field nulling, the sensors were operated in open loop so that the uniform component of the background field could be measured. The current is set by the PI controller providing updates at a 65 Hz frequency with *k*_*p*_ = 0.3*s* and *k*_*i*_ = 0.15*s* where *s* is the iteration step size used in static nulling. When dynamic field nulling was turned off, the current values were returned to the original values of the static field nulling. Due to limitations with the sensor firmware, closed loop operation was not possible during dynamic field nulling. However, we expect that the background field maintained by the nulling coils will minimize CAPE.

### 2.5. Source estimation of somatosensory evoked fields

To understand the impact of background fields on MEG data quality, we performed a median nerve stimulation (MNS) experiment which is a gold standard for functional mapping in epilepsy (Burgess et al., 2011). We repeated the recording three times in wearable MEG with dynamic field nulling, static field nulling, and no field nulling, and once in cryogenic MEG.

The protocol was approved by the Institutional Review Board at Massachusetts General Hospital (protocol #2025P000721). After providing informed consent and assent, a healthy 10-year-old child participated in our study. After placing electrodes connected to a constant current stimulator (model DS7A, Digitimer) over the median nerve, the stimulation intensity was adjusted until a visible thumb twitch was observed. 500 trials were performed with an inter-stimulus interval of 1.5 seconds. The experiment was repeated in wearable MEG (15 magnetometers) and cryogenic MEG (MEGIN Triux neo, 102 magnetometers). T1-weighted MRI was acquired separately for anatomical co-registration. For wearable MEG, the participant’s face was digitized during the recording with a 3D Camera (Creality Otter) which was later used for co-registration with the MRI using in-house software. For cryogenic MEG, digitization was performed using Polhemus Fastrak and used for MRI-MEG co-registration.

The wearable MEG data was bandpass filtered between 4 to 100 Hz using the default filter settings in MNE-Python (Gramfort et al., 2013). Notch filtering was performed at 60 Hz for line noise. After marking the bad sensor, homogenous field correction (HFC; (Tierney et al., 2021)) was applied with order = 1 to remove uniform field components from the data. The temporal Signal Space Separation (tSSS) algorithm was applied to the MEG data to remove artifacts due to background field (Taulu & Simola, 2006). The cryogenic MEG data was filtered the same way as wearable MEG data.

A boundary element model (BEM) was constructed using scalp surfaces extracted by Freesurfer and only magnetometers were used in the forward calculations. After extracting epochs around the events of interest, they were averaged to obtain event related fields (ERFs). The pre-stimulus baseline was used to compute a covariance matrix. Finally, equivalent current dipoles (ECDs) were computed at the time points of interest in the ERF (Sarvas, 1987).

## 3. Results

### 3.1 Dipole localization error of wearable MEG compared to cryogenic MEG

First, we compared the dipole localization error under different amounts of movement in wearable MEG vis-à-vis cryogenic MEG. The maximum rotations in each motion trajectory (**Fig. 3A**) are visualized in **Fig. 3B**. For small rotations, the dipole localization error in wearable MEG was smaller than cryogenic MEG since the simulated source location is superficial which allows favorable SNR for wearable MEG (**Fig 3C**)(Iivanainen et al., 2017). The dipole localization accuracy was best with no head movement (< 5 mm) and became worse with increasing amount of head movement. Cryogenic MEG did not allow motion trajectories beyond 12° maximum rotation due to the rigid gantry. Although wearable MEG allowed greater head movement than cryogenic MEG, with a 1 nT background field, the localization error in wearable MEG was almost 2 cm, too large for clinical use. Thus, while the 1 nT background field threshold determined by (Borna et al., 2022) is optimal for no head movement, it is not necessarily optimal for measurements with large head movements.

**Figure 3.**
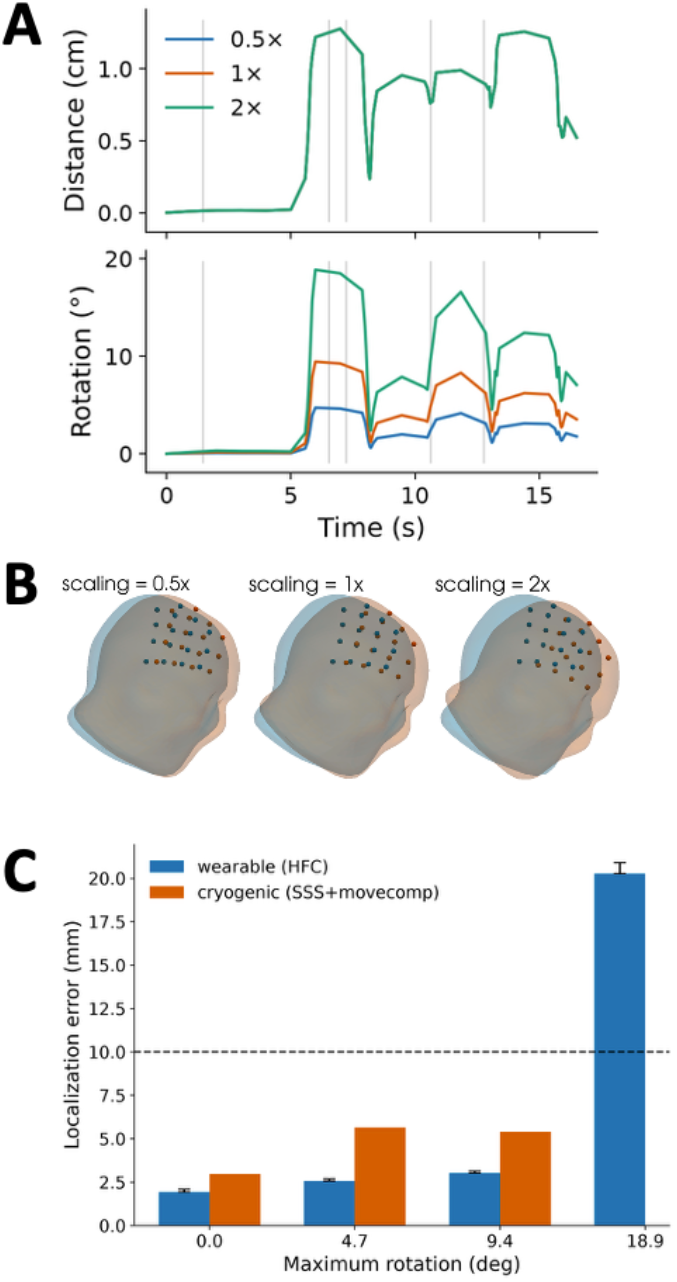
Comparison of dipole localization error in wearable MEG and fixed cryogenic MEG for 300 nAm interictal discharge and 1 nT background field. **A.** Motion trajectories with gray vertical lines showing times of simulated interictal discharges. The rotation in the original motion trajectory is scaled by a factor of 0.5 and 2 to test the impact of exaggerated motion. **B**. Visualization of the initial position and the position of maximum rotation in each of the three motion trajectories. **C**. Dipole localization error for no motion and the three motion trajectories.

### 3.2 Interaction of head motion and background field in wearable MEG

Next, we investigated the dipole localization error as a function of the background field amplitude for each of the movement trajectories. While the localization error was small for small background fields, it increased rapidly as the amount of background field increased. With residual background fields, the impact of head movement has a compounding effect on the localization error. Greater the background field, greater are the head movement artifacts and CAPE error. Thus, the tilt in the sensitive axes of the sensors degrades the effectiveness of spatial filtering methods more with larger background fields, where the movement artifacts are the largest. We determined the magnitude of the background field at which localization error remained acceptable (< 1 cm) for different sensor arrays. As the amount of head rotation increased, this background field threshold decreased (**Fig 4A**). The localization errors for wearable MEG in **Fig 3C** can be read from the y-axis values at 1 nT in **Fig 4A**. The dipole localization error rose abruptly beyond the background field threshold which underscores the importance of estimating it accurately. Interestingly, larger sensor arrays did not reduce the localization error (**Fig 4B**). This may be because a uniform background field introduced tilts to the sensitive axes in similar directions, resulting in an additive bias during dipole localization. In contrast, the simulations in (Borna et al., 2022) assumed a random background field direction per sensor, which would result in reduced variance with larger sensor arrays.

**Figure 4.**
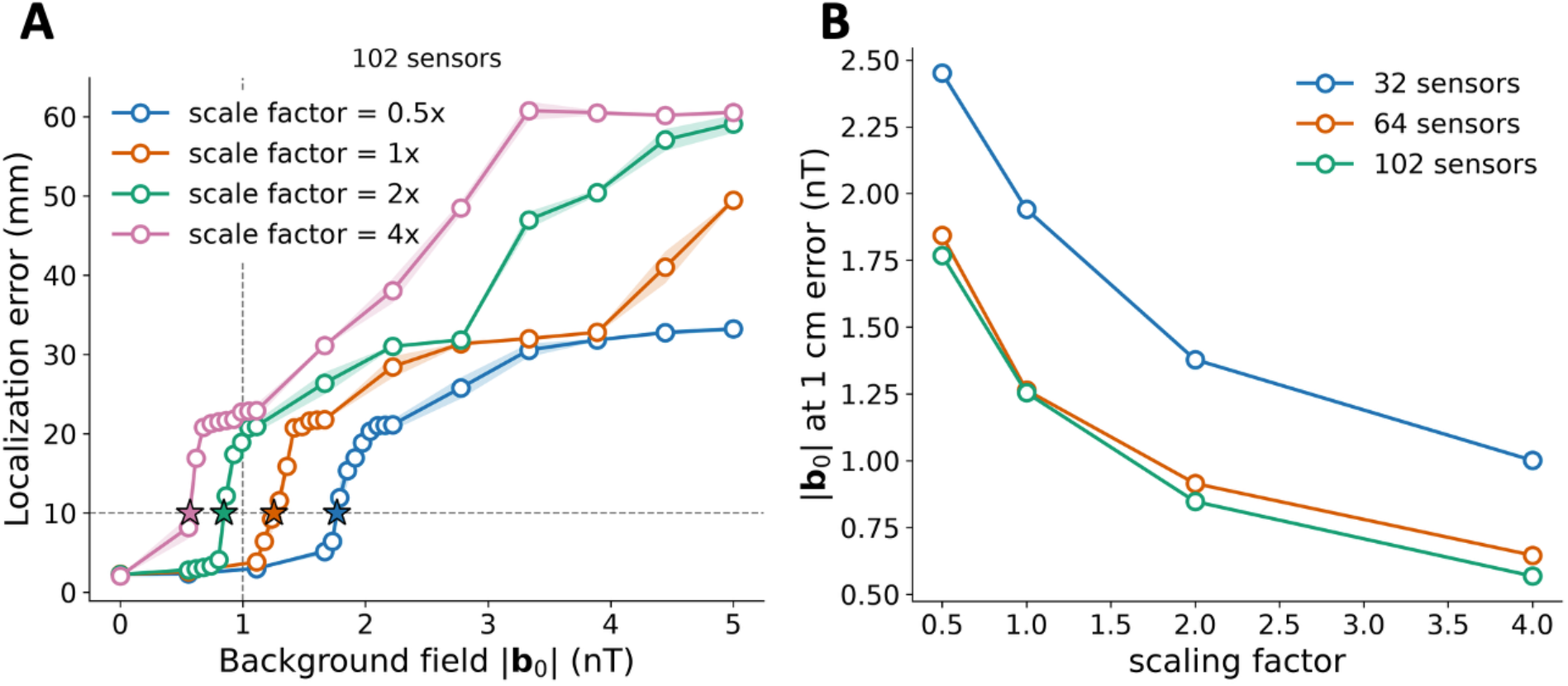
**A**. Mean dipole localization errors (with standard error of mean in shaded color) as a function of background field for different motion trajectories in a 102-sensor array. The background field threshold that achieved 1 cm localization error is shown with star sign and the 1 nT background field threshold is shown with a vertical line. **B**. The background field thresholds are plotted against the amount of motion for each of the sensor arrays.

### 3.5. Effectiveness of dynamic field nulling in improving source localization accuracy

Data acquired without any field nulling resulted in large background fields, sensor miscalibration and saturation that was not repaired even after application of homogenous field compensation (HFC) and high pass filtering (4 Hz). Due to CAPE-related miscalibrations in the sensor gain and sensitive axis, HFC cannot fully resolve the background field artifacts (**Fig 5A**). Static field nulling significantly mitigated these artifacts, but they were not fully resolved (**Fig 5B**). Dynamic field nulling in combination with HFC improved data quality by reducing both the magnitude and frequency of high-amplitude artifacts (**Fig 5C**). Note that the artifacts appear oscillatory due to ringing artifacts resulting from the data filtering. The mean background field in the reference sensors during dynamic field nulling (0.3 nT) was lower than the mean background field during static nulling (4.7 nT) and no nulling (21.3 nT). With dynamic field nulling, small imperceptible head movements did not result in significant artifacts.

**Figure 5.**
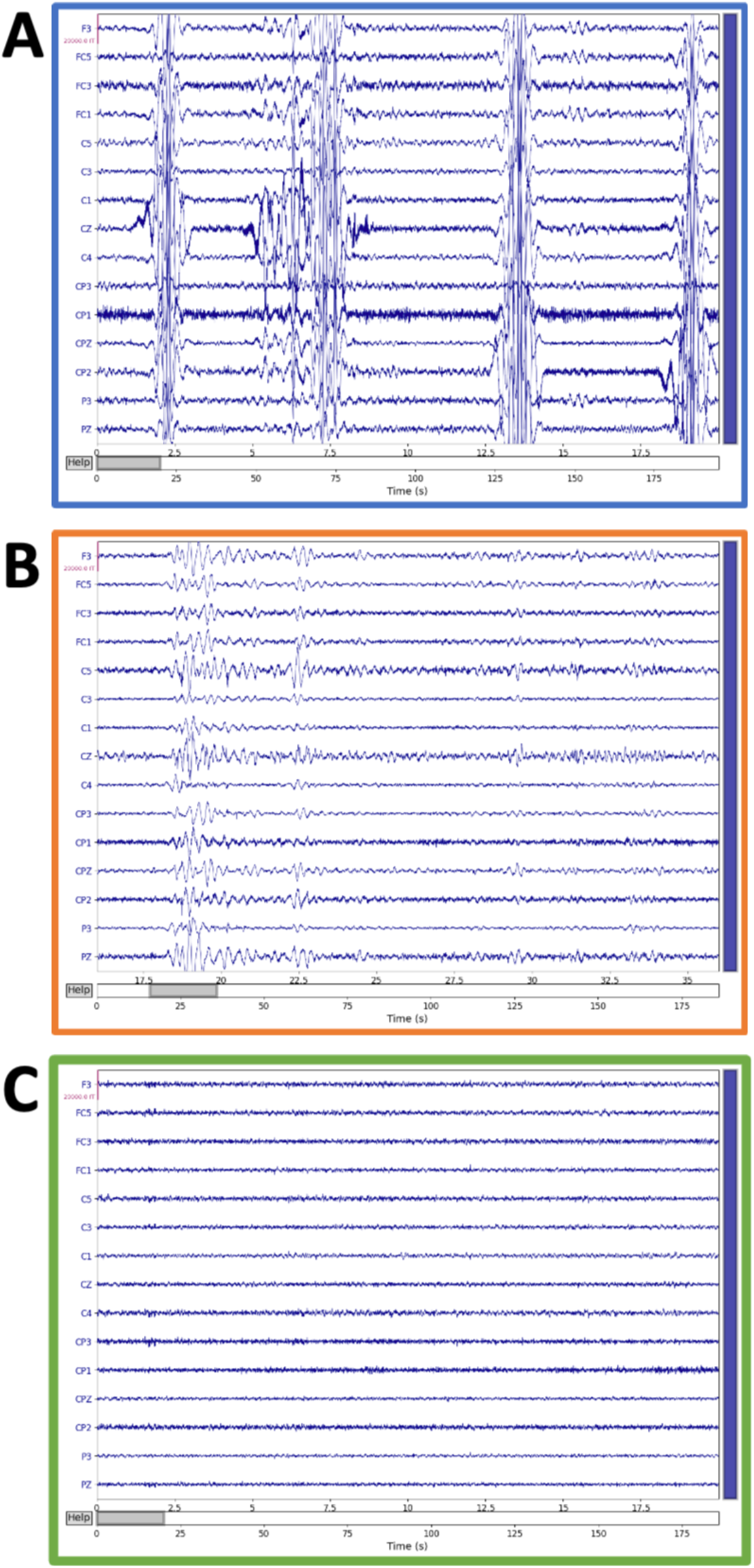
Representative 20 second data with (A) No field nulling, (B), static field nulling, and (C) dynamic field nulling. The data was processed with HFC and high-pass filtered at 4 Hz.

The artifacts due to background field impacted lower frequency components of the data (**Fig 6A**) more than higher frequencies. Application of static field nulling reduced these low-frequency artifacts, but dynamic field nulling achieved the best data quality. Dynamic nulling also improved the percentage of epochs retained after rejection (threshold 10 pT peak-to-peak) from 56% to 6% (**Fig 6B**).

**Figure 6.**
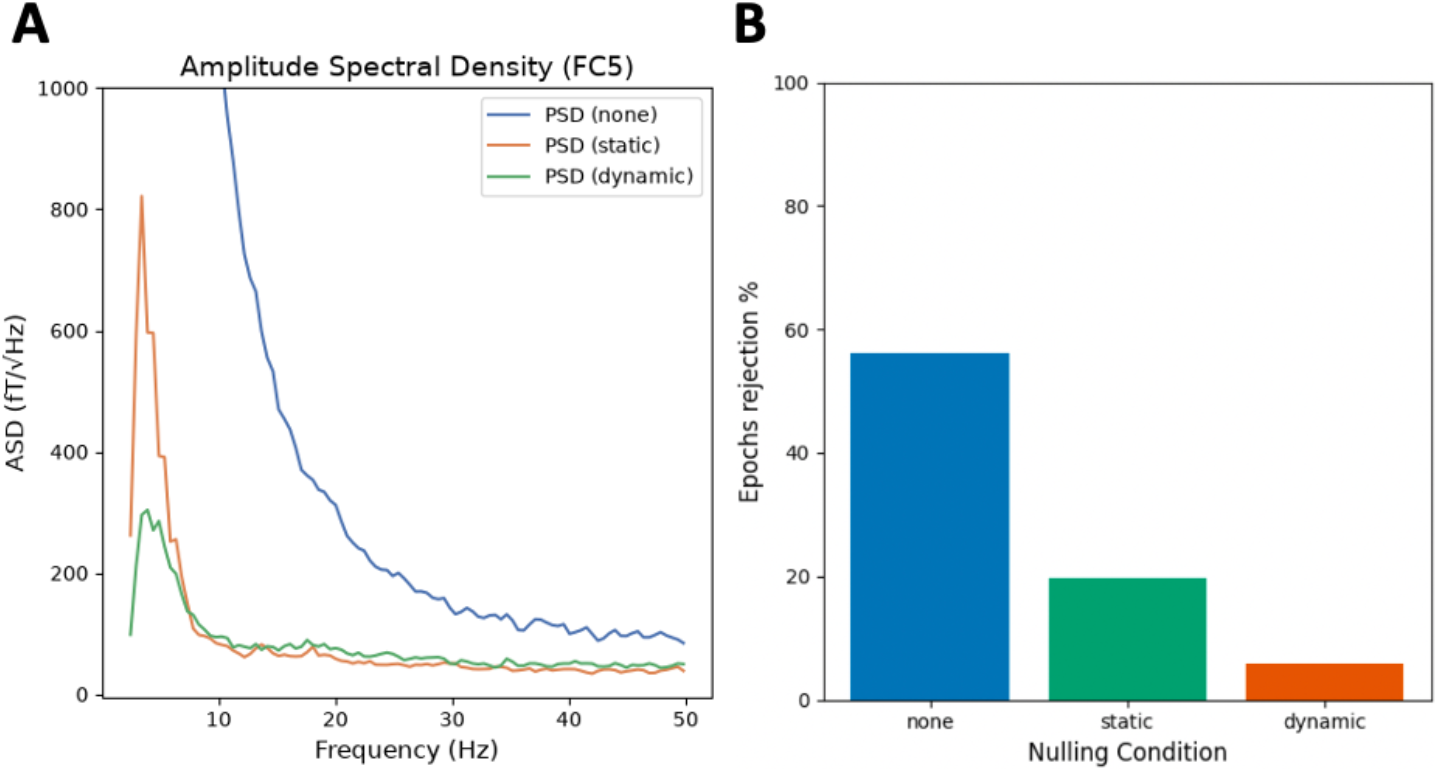
**A**. Amplitude spectral density of the processed continuous data with no field nulling (blue), static field nulling (green), and dynamic field nulling (orange). **B**. The percentage of epochs rejected with a 10 pT peak-to-peak threshold in the nulling conditions. Dynamic nulling retains a higher percentage of epochs than no field nulling.

The ERF closely matched cryogenic MEG with dynamic field nulling (**Fig 7A**) reproducing clinically relevant components including N20, P31, and P75. Dipole localization of these individual components (**Fig 7B, C**) revealed that dynamic nulling significantly improved accuracy compared to both no field nulling and static field nulling. Interestingly, static field nulling performed worse than no field nulling in terms of localization accuracy. This appears to be because static field nulling reduces the amplitude of the background-field-related motion artifacts below the data rejection threshold. Thus, the motion artifacts were not eliminated, but simply escape rejection, and instead corrupt the spatial topography of the measured brain activity.

**Figure 7.**
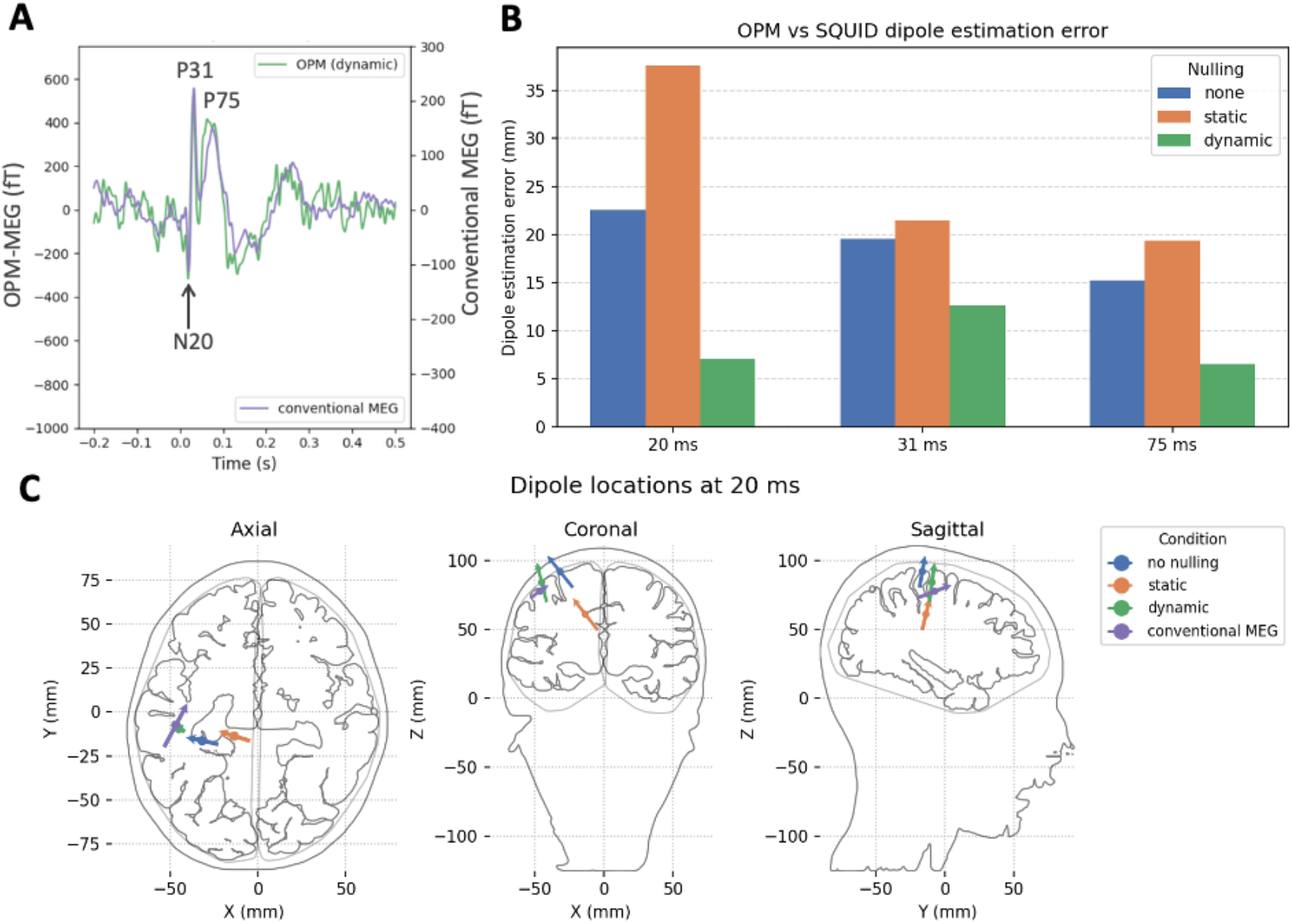
**A:** Event related field (ERF) from median nerve stimulation showing high concordance between conventional MEG (purple) and OPM-MEG (green) with dynamic field nulling. **B**. Dynamic field nulling improves accuracy of OPM-MEG dipole estimation at 20 ms (N20) from 2.2 cm without field nulling to 0.7 cm when compared to conventional MEG. **C**. The estimated dipole locations comparing conventional MEG (purple) to OPM-MEG with no nulling (blue), static nulling (orange) and dynamic nulling (green).

## 4. Discussion

We investigated the impact of head movement on dipole localization accuracy in wearable MEG. Our simulations demonstrate that although large head movements are possible in wearable MEG, they can adversely affect the dipole localization accuracy. To improve the dipole localization error, the background field had to be reduced using dynamic field nulling. In our wearable sensor array with 18 sensors (of which 3 are reference sensors), software compensation by itself was not adequate and it had to be combined with background field nulling to achieve more accurate localization. Our study highlights the importance of tracking the background fields in conjunction with the head motion for quality assurance in wearable MEG.

### Software-based artifact compensation complements but does not replace field nulling

Spatial filtering techniques can be used to estimate the spherical harmonic components of the background field and compensate for motion artifacts by zeroing them out (Tierney et al., 2021, 2024). However, to effectively separate the signal and noise subspaces, a larger number of sensors is often considered beneficial (Holmes et al., 2023; Tierney et al., 2022). Although this is true for perfectly calibrated sensor arrays, our simulations suggest that larger sensor arrays may be detrimental if the sensor arrays are not perfectly calibrated. While HFC can theoretically model and compensate both for uniform and gradient components of the background fields, in our experience, its effectiveness in mitigating field gradients was limited for small sensor arrays. Finally, hardware-based compensation reduces the magnitude of motion artifacts and the associated complexity of analyzing such data, often a hurdle for clinicians and neuroscientists. Thus, software and hardware compensation of background fields are complementary methods of mitigating motion artifacts.

### On-sensor field nulling mitigates CAPE errors but not motion artifacts

On-sensor field nulling can mitigate CAPE errors but without reducing the background field in a region around the head, motion artifacts are still expected. Unless the CAPE error is completely eliminated, a fraction of these large artifacts will persist even after spatial filtering is applied and result in dipole localization errors. As research-grade wearable MEG approaches dense sensor layouts to fully exploit the available on-scalp signals, crosstalk due to on-sensor field nulling is an increasing concern. PCB-based background field nulling will reduce these on-sensor cancellation currents and mitigate the need to model such crosstalk effects (Nardelli et al., 2019).

### Sensor design for mitigating motion artifacts

Triaxial sensors measure tangential field components in addition to the radial field components and the additional channels are considered advantageous for suppression of motion artifacts compared to single or dual-axis sensors (Brookes et al., 2021). Gradiometers are another design that can remove background fields by removing common mode noise (M. Hämäläinen et al., 1993b). Nonetheless, both triaxial sensors and gradiometers increase the cost of MEG sensor arrays with gradiometers requiring additional vapor cells. On the other hand, background field nulling using PCB-based coils is a cost-effective solution that requires only a single reference triaxial sensor.

### Modeling limitations

We did not model gradient fields in our current work. Gradient fields are inherently more complex to model and null due to concomitant field components which require adequate sampling with reference sensors. The gradient field depends on the dimensions of the shielded room and characteristics of nearby sources of magnetic fields. Our prior publication also suggested that the gradient fields in our shielded room are potentially small (Jas et al., 2025). While uncompensated uniform fields can result in artifacts primarily due to sensor rotation, uncompensated gradient fields can result in artifacts even during linear translation of the sensors. The experimental data suggests that combining the hardware compensation with HFC is sufficient in our magnetic field environment. Future work is needed to determine the thresholds for background field gradients and to mitigate artifacts due to them. Ultimately, the concurrence of brain activity and movement presents the greatest challenge and opportunity in wearable MEG.

## 5. Conclusion

The background field thresholds can provide practical guidelines for clinicians using wearable MEG in diagnosis and epilepsy surgery planning. In epilepsy patients, particularly children who move frequently, tracking of both head motion and background field can be critical for improving reliability and interpretability of the dipole localization. Although visual inspection of data is crucial, motion artifacts with field amplitudes comparable to brain activity can be particularly deceptive. As our analysis of static field compensation suggests, partial compensation of background field can be potentially worse than no compensation as it may provide misleading results during data analysis. By discarding data segments with excessive motion, we can improve reliability of dipole localization. At the same time, our work underscores the importance of developing systems that track head motion relative to the background field. While in cryogenic MEG, head motion is tracked relative to the sensor array, wearable MEG requires head motion to be tracked relative to the background field. The advantage of our PCB-based field nulling coils is that their precise geometry will allow head motion tracking in addition to background field nulling (Iivanainen et al., 2022). Ultimately, our work emphasizes the need for MEG setups with tightly integrated hardware and software.

## 6. Data and Code Availability

The data and code will be made available on GitHub and OSF upon publication.

## 7. Author Contributions

**Mainak Jas:** Conceptualization, Methodology, Software, Validation, Investigation, Resources, Writing – original draft, Writing – review & editing, Visualization, Project administration, Funding acquisition. **Teppei Matsubara:** Methodology, Validation, Investigation, Visualization, Writing – review & editing. **Steven M. Stufflebeam:** Conceptualization, Resources, Writing – review & editing. **Padmavathi Sundaram:** Methodology, Investigation, Writing – review & editing. **Seppo P. Ahlfors:** Validation, Investigation, Visualization, Supervision, Writing – review & editing.

## 8. Funding

This study was supported by NIH grants R21NS140619, S10OD030469, and P41EB030006. The content is solely the responsibility of the authors and does not necessarily represent the official views of the National Institutes of Health.

## 9. Declaration of Competing Interests

Steven M Stufflebeam is a co-founder at FIND Neuro. The remaining authors declare no competing interests.

## Notes

### Summary of Updates

Figure 3 and Figure 4 updated and the corresponding text clarified and updated.

